# Imputed graph-genotyped structural variants identify regulatory haplotypes associated with gene expression in Atlantic salmon

**DOI:** 10.64898/2026.06.03.729980

**Authors:** Maëlys Chapis, Domniki Manousi, Célian Diblasi, Cathrine Brekke, Jun Soung Kwak, Arturo Vera Ponce De Leon, Mariann Arnyasi, Randi Fenstad, Solomon Antwi Boison, Marie Saitou

**Affiliations:** Université de Bretagne Occidentale, Brest, France; Department of Animal and Aquacultural Sciences, Faculty of Biosciences, Norwegian University of Life Sciences, Ås, Norway; The Roslin Institute and Royal (Dick) School of Veterinary Studies, University of Edinburgh, Easter Bush, Midlothian, United Kingdom; Department of Biology, Lund University, Lund, Sweden; Department of Aquatic Life Medicine, Kongju National University, Chungcheongnam-do, South Korea; Mowi Genetics AS, Norway

**Keywords:** structural variants, eQTL, recombination rate, gill tissue, graph genotyping, genotype imputation, regulatory haplotypes

## Abstract

Structural variants (SVs) can affect gene regulation, but they are difficult to include in expression genetic studies when large RNA-seq cohorts lack whole-genome sequencing. This is common in non-human and non-model systems, where whole-genome sequencing at population scale remains costly. As a result, expression quantitative trait locus (eQTL) studies often rely on single nucleotide polymorphism (SNP) markers. These analyses can identify expression-associated regions, but often provide limited biological interpretation of the underlying regulatory mechanisms. Here, we used Atlantic salmon as a study system to test whether graph-genotyped SVs can be imputed into a SNP-array-genotyped RNA-seq cohort and used to interpret regulatory haplotypes. SVs were discovered from two long-read-sequenced individuals, supplemented with short-read SV and SNP calls from a 112-individual whole-genome-sequenced reference panel, graph-genotyped, jointly phased with SNPs, and imputed into 906 offspring with gill RNA-seq and SNP-array genotypes. After size filtering, the imputed SV catalogue contained 100,269 variants and showed nonuniform genomic distributions associated with sex-specific recombination landscapes. Association testing identified 51 SV-eQTL candidates, including 35 cis and 16 trans associations. These candidates were enriched for short-read-derived variants, indicating that short-read supplementation can recover regulatory variants missed by small-scale long-read discovery. SV-eQTL candidates were more strongly tagged by nearby SNPs than non-associated variants generally, but individual SNP lead markers often failed to capture the same eQTL signals in conditional regression. Retained candidates after the conditional analysis included target-gene-overlapping deletions, nearby local variants without target-gene overlap, trans associations, and short insertions with opposite effects on gene expression. These results show that imputed graph-genotyped SVs can add biological interpretation to possible regulatory haplotypes.

## Introduction

Gene expression variation provides a molecular link between genetic variation and phenotypic diversity. Differences in transcript abundance can alter cellular state, developmental trajectories, immune responses, metabolic activity, and other organismal processes, thereby providing one route through which DNA sequence variation can influence traits. Expression quantitative trait locus (eQTL) mapping is used to identify genomic loci at which genetic differences among individuals are associated with differences in gene expression. In a typical eQTL analysis, gene expression is treated as a quantitative phenotype, and nearby or genome-wide genetic markers are tested for association with transcript abundance. Large-scale eQTL resources, including studies with hundreds to thousands of individuals across many tissues, have shown that genetic effects on expression are widespread and can help connect noncoding genetic variation to molecular phenotypes (GTEx Consortium 2020; Scott et al. 2021; Mostafavi et al. 2023; Wang et al. 2024; Bian et al. 2025; Brotman et al. 2025; Renganaath and Albert 2025; Li et al. 2026).

However, the lead SNP at an eQTL, usually defined as the SNP with the strongest statistical association with expression at a locus, does not necessarily identify the mechanism underlying the expression difference (Connally et al. 2022). In many cases, the lead SNP is better understood as a marker of an associated regulatory haplotype (Mostafavi et al. 2023; Reales et al. 2026): it indicates that expression-associated genetic variation is likely to reside in that genomic neighborhood, but it does not by itself specify which sequence feature changes expression. This distinction is important because SNP-based eQTL mapping can localize regulatory variation to a region or haplotype, while leaving unresolved whether the relevant molecular allele is a SNP, an indel, a repeat, a structural variant, or another linked feature. Thus, SNP-based eQTL analysis provides a powerful way to find genomic regions associated with expression variation, but additional information is needed to move from locating regulatory signals to interpreting the causal variants that underlie them (Funk et al. 2025; Mudappathi et al. 2025).

Genomic structural variants (SVs) are a particularly important class of variants to examine after SNP-based eQTL mapping because they alter genomic sequence at scales and in configurations that can be more directly connected to molecular function than individual SNPs (Hämälä et al. 2021; Fang and Edwards 2024; Stuart et al. 2026). Deletions, insertions, duplications, inversions, and other rearrangements can alter gene dosage, transcript structure, untranslated regions, promoters, enhancers, repeat-associated regulatory elements, or local chromatin organization (Karageorgiou et al. 2026). In this context, an SV can provide a concrete structural hypothesis for an expression-associated haplotype: the haplotype is no longer represented only by an anonymous marker SNP, but by a specific deletion, insertion, duplication, inversion, or other structural variant.

Several studies have established that SVs contribute to regulatory variation and should not be excluded from eQTL interpretation. In humans, early genome-wide SV-eQTL work showed that SVs can act as lead regulatory markers and contribute to expression variation (Chiang et al. 2017). Subsequent analyses of SVs and short tandem repeats in pluripotent stem cells showed that these variant classes can influence gene expression, differ in their regulatory properties, and intersect complex trait-associated loci (Jakubosky et al. 2020). Deep whole-genome sequencing in GTEx further showed that common SVs are enriched among putative causal eQTL variants and can affect the expression of multiple nearby genes, emphasizing that SVs can provide regulatory information not captured by SNP markers alone (Scott et al. 2021). Multi-omic studies have extended this view by linking SVs to expression, splicing, chromatin modification, and protein abundance, indicating that SV effects can propagate across several molecular layers (Vialle et al. 2022).

SV-associated regulatory effects have also been reported in humans (Chiang et al. 2017; Jakubosky et al. 2020; Scott et al. 2021; Kirsche et al. 2023; Bai et al. 2026 May 20), in other animals, including livestock systems (Bhati et al. 2023; Falker-Gieske et al. 2023; Leonard et al. 2023; Leonard et al. 2024; Yang et al. 2024), and in plants (Zhou et al. 2022; Cao et al. 2024; Li et al. 2024; Zhang et al. 2024; Yildiz et al. 2025), where population-scale SV resources have been linked to gene expression, regulatory divergence, and trait variation. Some of such studies have shown that regulatory loci can contain multiple linked or partially linked signals, and SV-eQTL studies have shown that SVs can be incompletely tagged by nearby SNPs and can affect more than one gene (Taylor et al. 2024). These studies motivate examining SVs not only as possible causal variants, but also as potential components of regulatory haplotypes. Under this view, an SV-eQTL may reflect an SV allele that is carried on, and potentially contributes to, an expression-associated haplotype whose effects can be evaluated alongside SNP markers.

Existing SV-eQTL studies have established that structural variants can contribute to regulatory variation, but many of these studies have relied on whole-genome sequencing or direct SV genotyping in the same individuals used for expression profiling. This design enables direct testing of SV-expression associations and has been highly informative in humans and other well-resourced systems, including studies that integrate SVs with RNA-seq, splicing, chromatin, or other molecular phenotypes (Bai et al. 2026 May 20). However, many non-model species do not possess such population-scale omics data sets. Whole-genome sequencing at sufficient depth for reliable variant discovery and genotyping remains costly for large cohorts, especially when long-read sequencing is required to resolve structural variants. By contrast, SNP arrays and RNA-seq are often more affordable as they generate smaller data volume, making them more commonly available in non-model populations. As a result, large RNA-seq cohorts may be accompanied by SNP-array genotypes, pedigree information, and occasionally smaller WGS reference panels. This creates a scale mismatch between transcriptomic phenotype data, which can be measured in many individuals, and structural variants genotypes, which are often observed directly only in smaller discovery or reference panels.

Graph genotyping and haplotype-based imputation provide a practical route for addressing this scale mismatch (Long et al. 2022; Meisner and Albrechtsen 2022; Yang et al. 2025; Sun and Li 2026). In graph genotyping, SVs are first discovered in a limited number of individuals using long-read sequencing, and the resulting SV catalogue is then genotyped in a larger short-read WGS reference panel using graph-based methods. Graph genotyping allows reads from each reference individual to be compared against alternative alleles, producing SV genotypes for the reference panel. These SV genotypes can then be phased together with SNPs to define joint SNP-SV reference haplotypes. In individuals with SNP-array genotypes but no WGS, haplotype-based imputation can use the SNP background to project SV genotypes from the reference panel into the larger cohort. Thus, graph genotyping and imputation can bridge small SV-observed reference panels and large SNP-genotyped expression cohorts, while preserving the distinction between directly observed SV genotypes and imputed haplotype labels.

However, this technical bridge has rarely been connected to RNA-seq-based regulatory interpretation in a single analysis. Existing studies have most often followed one of two designs. One line of work has directly mapped SV-eQTLs in cohorts with matched WGS and transcriptomic or multi-omic data, establishing that SVs can contribute to expression variation, splicing, chromatin state, protein abundance, and multi-gene regulatory effects (Chiang et al. 2017; Jakubosky et al. 2020; Scott et al. 2021; Vialle et al. 2022). Another line of work has focused on graph-based genotyping or imputation of SVs into larger cohorts, often for population-scale variant representation, GWAS, or trait association rather than RNA-seq-based regulatory interpretation (Eggertsson et al. 2019; Long et al. 2022; Meisner and Albrechtsen 2022). Fewer studies have connected SV discovery, graph-based SV genotyping, haplotype-based imputation, RNA-seq-based eQTL mapping, SNP-eQTL comparison, and haplotype interpretation within the same framework. As a result, it remains unclear how graph-genotyped SVs, once projected into a SNP-genotyped expression cohort, should be interpreted relative to SNP-eQTL lead markers and local tagging SNPs. A suitable test case for this problem requires two conditions: a genome in which structural variant representation is nontrivial, and a population resource in which SNP genotypes, pedigree information, WGS reference individuals, and RNA-seq phenotypes can be connected.

Atlantic salmon provides an informative setting for evaluating this integrated framework because its genomic resources have developed through several complementary but only partly connected phases. Early genome studies established Atlantic salmon as a structurally complex vertebrate genome shaped by the salmonid-specific whole-genome duplication, subsequent rediploidization, extensive duplicated sequence, repeat-associated complexity, and large-scale genome restructuring (Lien et al. 2016; Sinclair-Waters et al. 2020; Gundappa et al. 2022). Dense SNP-array resources and pedigree-based imputation, based on the genomic resources, then enabled large-scale genomic analyses in breeding populations, including trait mapping and genomic prediction using increasingly high-density SNP information (Houston et al. 2014; Yáñez et al. 2016; Tsai et al. 2017; Sinclair-Waters et al. 2020; Tsairidou et al. 2020; Sinclair-Waters et al. 2022; Gundappa et al. 2025). In parallel, population-scale WGS and long-read resources began to reveal the structural-variant landscape of Atlantic salmon, including curated SV catalogues, inversion polymorphisms, and SV differentiation among populations (Bertolotti et al. 2020; Stenløkk 2023; Lecomte et al. 2024). More recently, SNP- and SV- imputation and small-sample multi-omics QTL studies have shown that both structural variants and molecular regulatory phenotypes can be analysed in this species, including a 12-individual multi-tissue dataset with matched WGS, RNA-seq, and ATAC-seq layers (Arnesen 2025; Nguyen et al. 2025; Macqueen et al. 2026).

Building on these resources, we used Atlantic salmon to test three linked predictions about the role of structural variants in expression-associated haplotypes. First, we expected that imputed SVs projected into the RNA-seq cohort would retain genome-architecture signatures of structural variation, including nonuniform distributions across chromosomes and recombination landscapes. Second, we expected that a subset of imputed SVs would be associated with gene expression and that these SV-eQTL candidates would not simply mirror the full imputed SV catalogue, but would be enriched for particular genomic contexts or discovery sources. Third, we expected that SV-eQTL signals would be only partially represented by nearby SNP markers: if imputed SVs provide structural interpretation of regulatory haplotypes, then some SV effects should persist after conditioning on SNP-eQTL lead markers or local SNP tags, while others should be attenuated by strong SNP tagging. We addressed these predictions by linking three layers of information that are generally analysed separately: SV alleles defined by graph-based genotyping in a WGS reference panel, SNP-defined haplotypes in a large offspring cohort, and gene expression variation measured by RNA-seq.

## Materials and Methods

### Study design and datasets

This study was designed to evaluate whether graph-genotyped structural variants (SVs) can be imputed into a SNP-array-genotyped RNA-seq cohort and used to interpret regulatory haplotypes associated with gene expression. The workflow connected three data layers: SV discovery and graph-based genotyping in a whole-genome-sequenced reference panel, haplotype-based imputation into an offspring expression cohort, and SV-eQTL analysis using gill RNA-seq expression phenotypes (**Figure 1**).

**Figure 1.**
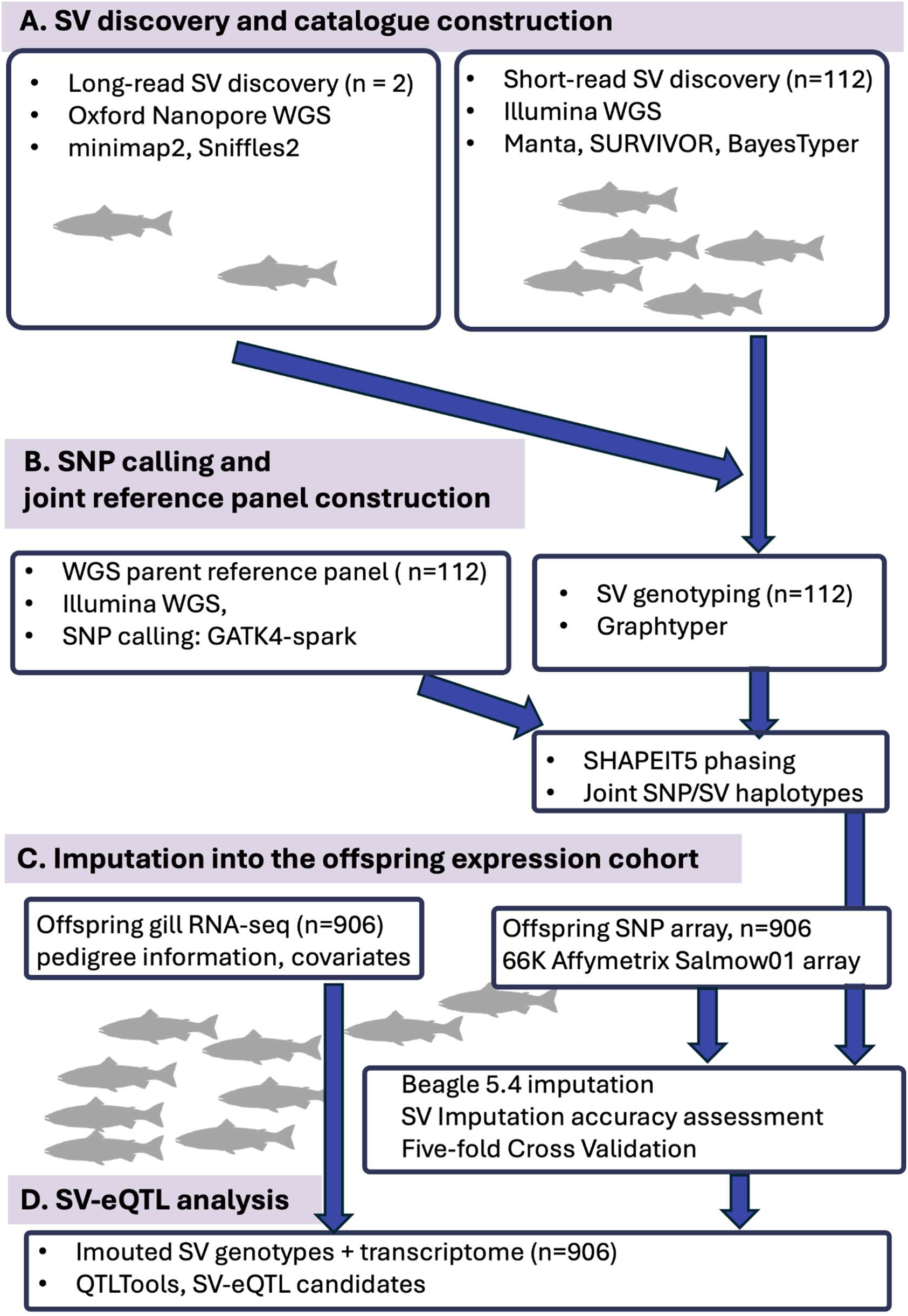
Study design and analytical workflow for imputing graph-genotyped structural variants into an Atlantic salmon expression cohort. **A. SV discovery and graph-based genotyping.** Structural variants (SVs) were discovered from two Oxford Nanopore long-read-sequenced individuals using Sniffles2 and supplemented with short-read-derived SV calls from 112 whole-genome-sequenced reference individuals. The combined SV catalogue was genotyped in the WGS reference panel using GraphTyper. **B. SNP calling and joint reference haplotype construction.** SNPs were called from the same 112 WGS reference individuals and combined with graph-genotyped SVs to construct a joint SNP/SV reference panel. Reference genotypes and offspring SNP-array genotypes were phased with SHAPEIT5, incorporating pedigree information where applicable. **C. Imputation into the offspring expression cohort.** SV dosages were imputed with Beagle 5.4 into 906 offspring with SNP-array genotypes and gill RNA-seq data. Imputation accuracy was assessed by five-fold cross-validation within the WGS reference panel. **D. SV-eQTL analysis and conditional analysis with local SNPs** Imputed SV dosages were tested for association with gill gene expression using QTLtools. SV-eQTL candidates were then analysed for genomic context, gene overlap, local SNP linkage disequilibrium, comparison with SNP-eQTL lead SNPs, and conditional retention after accounting for either SNP-eQTL lead SNPs or the highest-LD nearby SNP.

Three connected datasets were used (See **Data Availability** section). First, long-read sequencing data were generated from two Atlantic salmon individuals from a Mowi breeding nucleus and used for initial SV discovery. Second, whole-genome sequencing (WGS) data from 112 related or ancestral individuals from the same breeding population were used as a reference panel for SNP and SV genotyping. Third, 906 offspring with SNP-array genotypes and gill RNA-seq data were used as the target expression cohort. SVs genotyped in the WGS reference panel were projected into the offspring cohort by haplotype-based imputation and then tested for association with gene expression.

All coordinate-based analyses used the Atlantic salmon reference genome Ssal_v3.1 / GCA_905237065.2.

### Long-read sequencing and initial SV discovery

Two Atlantic salmon individuals, derived from a breeding nucleus of an aquaculture company (MOWI), were used for long-read sequencing. High molecular weight genomic DNA was extracted from blood samples using the Monarch HMW DNA Extraction Kit for blood (New England Biolabs, USA), following the manufacturer’s protocol. For each individual, 5 µg of purified genomic DNA was obtained. The genomic DNA was mechanically fragmented using needle shearing (10 passes through a 21-gauge needle) to reduce viscosity and improve compatibility with ONT Flongle sequencing. Size reduction was confirmed by agarose gel electrophoresis. Sequencing libraries were prepared using the Ultra-Long DNA Sequencing Kit V14 (Oxford Nanopore Technologies, UK) according to the manufacturer’s instructions. The PromethION flow cell (Oxford Nanopore Technologies, UK) was used for sequencing. Flow cell maintenance and cleaning were performed using the Flow Cell Wash Kit (Oxford Nanopore Technologies, UK).

Basecalling was performed using Dorado (version v0.5.0]) with default parameters. The resulting reads were quality filtered using Filtlong v0.2.1 to remove the worst 10% of total reads as well as reads with length shorter than 4000 bp. Filtered reads were then aligned to the Atlantic salmon reference genome (Ssal_v3.1, GCA_905237065.2) using minimap2 (Li 2018). Oxford Nanopore sequencing of the two Atlantic salmon individuals generated 3.65 million quality-filtered reads, corresponding to 89.43 Gb of sequence data. Per-sample sequencing yields were 54.24 Gb for one and 35.19 Gb for another. Mean read lengths were 26.25 kb and 22.28 kb, respectively, with read N50 values of 47.56 kb and 42.11 kb. The maximum read lengths were 368.99 kb for TT20 and 445.78 kb for CC03. The GC content was similar between samples, at 43.47% and 43.45%, respectively.

### Short-read SV supplementation and graph-based SV genotyping

Structural variants (SVs) were identified in each individual using Sniffles2, and subsequently merged to establish an initial SV catalog (Smolka et al. 2024). To supplement this catalogue, we incorporated SVs detected from Illumina based WGS data from the 112 reference individuals using Manta (Diblasi et al. 2026). These individuals were close relatives of the two long-read individuals in Mowi breeding stock. Short-read data were quality filtered with fastp v0.23.2 (https://github.com/opengene/fastp) and aligned to the Atlantic salmon reference genome Ssal_v3.1 using bwa-mem2 (Vasimuddin et al. 2019).

The short-read-derived SVs had initially been genotyped with BayesTyper, and the combined long-read and short-read SV catalog was subsequently genotyped in the 112 WGS individuals using GraphTyper v2.7.5 (Eggertsson et al. 2019). The final catalog therefore included both long-read-detected SVs and short-read-detected SVs. GraphTyper output included symbolic SV alleles as well as sequence-resolved variants; therefore, downstream summaries distinguished between the full imputed variant set, the VEP-annotated variant set, and size-filtered SVs. Genotyped SVs were filtered using PLINK2 by removing variants with more than 20% missing genotypes or minor allele frequency below 2%.

### SNP calling in the WGS reference panel

SNP genotypes for the WGS reference individuals were obtained from a previously described Atlantic salmon WGS dataset (Buso et al. 2025). Briefly, Illumina reads were mapped to the Atlantic salmon reference genome Ssal_v3.1 (GCA_905237065.2) using bwa-mem2. Duplicate reads were marked with GATK4 Spark MarkDuplicates, variants were called with GATK4 Spark HaplotypeCaller, and individual genotypes were merged with GATK4 Spark GenotypeGVCFs. SNPs were hard-filtered using standard GATK criteria for quality by depth, variant quality, strand bias, and read-position bias. Biallelic SNPs were then retained using VCFtools (Danecek et al. 2011) with depth >4 and <30, genotype quality call rate >90%, and missingness <10%. The filtered SNPs from the 112 Mowi WGS individuals were used as the SNP component of the reference panel for haplotype phasing and imputation.

Genotyped SNPs and SVs for the 112 WGS individuals were filtered using PLINK2 (Chang et al. 2015) and different filtering criteria based on the different types of genetic variation. In particular, SVs with more than 20% missing genotypes and minor allele frequency (MAF) below 2% were removed, whereas SNPs were filtered based on Hardy-Weinberg Equilibrium (HWE p-value < 1E-8), minor allele frequency (MAF < 0.01), and genotype missingness thresholds per individual and SNP variant (10% and 5%, respectively). The filtered WGS SNPs were used together with graph-genotyped SVs to construct the joint SNP/SV reference haplotype panel.

### Offspring SNP-array genotypes and RNA-seq expression data

The present analysis used the continuous-light subset of the broader RNA-seq experiment because this subset had matched SNP-array genotypes, gill RNA-seq expression data, and pedigree structure suitable for haplotype-based imputation. The original RNA-seq experiment included 2,970 fish reared under three photoperiod conditions. In the present study, we used the subset of 906 fish from the continuous-light (LL) condition. A detailed description of the sampling and generation of the transcriptomics information of these fish is provided elsewhere (Gjerde et al. 2025; Saitou et al. 2025).

SNP-array genotyping was performed using the 66K Affymetrix SNP array Salmow01. Physical array coordinates were reassigned to the Atlantic salmon reference genome Ssal_v3.1 (GCA_905237065.2), and genotypes were quality-filtered using PLINK v1.9 (Purcell et al. 2007). Variants were removed if they had Hardy-Weinberg equilibrium P < 1 × 10⁻⁶, MAF < 0.01, or missing genotype calls in more than 3% of individuals. Samples with more than 10% missing genotypes were removed. After filtering, 48,563 SNP-array markers were retained.

The SNP-based imputed genotype set used for SNP-eQTL comparison had been generated previously for the same offspring cohort. Briefly, SNP genotype imputation was performed using Beagle 5.2 (Browning et al. 2018; Browning et al. 2021) with WGS parent genotypes as the reference panel. Imputation accuracy was evaluated by five-fold cross-validation in the parent fish, masking genotypes in 20% of individuals and imputing them from the remaining 80%. Accuracy was calculated as the squared Pearson correlation coefficient (R²) between true and imputed genotypes, averaged across validation folds. This procedure retained 412,069 imputed SNPs with R² > 0.80, which were further filtered for Hardy-Weinberg equilibrium using PLINK v1.9 with a threshold of P < 1 × 10⁻⁸.

RNA-seq data generation and expression quantification were described previously by Grønvold et al. Briefly, gill biopsies were flash frozen, RNA was extracted using the RNeasy Fibrous Tissue Mini Kit, and libraries were sequenced using 2 × 150 bp paired-end Illumina sequencing. Adaptor sequences were removed with fastp v0.23.2. Transcript abundance was quantified with Salmon v1.1.0 (Patro et al. 2017) using the Atlantic salmon transcriptome annotation from Ssal_v3.1. Selective mapping was performed using a transcriptome index, and the --keepDuplicates and --gcBias options were used to account for high similarity among duplicated genes and fragment-level GC bias, respectively. Gene-level expression was calculated by summing transcript-level raw counts and normalizing to transcripts per million using tximport.

### Ethics statement

The original RNA-seq experiment included 2,970 fish reared under three photoperiod conditions. In the present study, we used the subset of 906 fish from the continuous-light (LL) condition because this group had matched SNP-array genotypes, gill RNA-seq data, and the pedigree structure required for haplotype-based imputation and eQTL analysis. The broader animal experiment from which the offspring RNA-seq and genotype data were derived was performed according to EU regulations concerning the protection of animals used for scientific purposes (Directive 2010/63/EU). Appropriate measures were taken to minimize pain and discomfort. The experiment was approved by the Norwegian Food Safety Authority (FOTS ID 25658). The studies were conducted in accordance with local legislation and institutional requirements, and permission to use the animals and associated samples was obtained from the owner. The fish material used for long-read sequencing was obtained from Atlantic salmon sampled during routine slaughter at Mowi. No additional experimental procedures were performed on these fish for the purpose of this study.

### Joint SNP/SV phasing and SV imputation into the offspring cohort

Graph-genotyped SVs and WGS-derived SNPs from the 112 reference individuals were combined to construct a joint SNP/SV reference panel. Allele conformation between overlapping WGS and SNP-array variants was checked using conform-gt. Reference and target genotypes were phased using SHAPEIT5 (Hofmeister et al. 2023). Pedigree information was incorporated where applicable to improve phasing accuracy in the offspring cohort. SV dosages were imputed into the 906 offspring using Beagle 5.4(Browning et al. 2018; Browning et al. 2021), with the 112 WGS individuals serving as the reference panel and the SNP-array-genotyped offspring serving as the target cohort. The resulting imputed SV dosages were used for SV-eQTL mapping, local SNP tagging analyses, and conditional regression analyses. SV imputation accuracy was evaluated by five-fold cross-validation within the WGS reference panel. In each fold, SV genotypes were masked in a subset of reference individuals and imputed using the remaining individuals as the reference. Imputed and observed genotypes were compared using the squared Pearson correlation coefficient, reported as imputation R². SV-eQTL candidates were retained only if their average imputation R² exceeded 0.5.

### SV catalogue characterization and genome-wide distribution analyses

SVs were characterized by variant type, size, genomic position, discovery source, and annotation overlap. Variant classes included deletions, insertions, duplications, inversions, and other GraphTyper-represented alleles where applicable. Size-based summaries were performed separately from full-catalogue summaries because some GraphTyper-represented alleles included variants at or below the conventional SV size threshold.

SV density was summarized in 1 Mb genomic windows across the Atlantic salmon genome. To examine the relationship between SV density and genome structure, SV counts per window were compared with gene density and sex-specific recombination-rate estimates from Atlantic salmon aquaculture populations (Brekke et al. 2023). Female and male recombination maps were analysed separately. Windows were assigned to recombination-rate quartiles within each sex, and SV density was compared among quartiles using Kruskal-Wallis tests. Chromosome-wise Spearman correlations were calculated between SV density and recombination rate separately for female and male maps.

Allele-frequency distributions were compared between long-read-derived and short-read-derived SVs retained in the imputed dataset. Minor allele frequency distributions were compared using Wilcoxon rank-sum tests and Kolmogorov-Smirnov tests. Rare and common allele-frequency categories were summarized descriptively.

To evaluate discovery-panel ascertainment, a simple diploid sampling model was used. For an allele with population frequency p, the probability of observing at least one alternate allele in a discovery panel of N diploid individuals was calculated as:

P = 1 - (1 - p)^(2N)

This model was used to estimate the number of diploid individuals required to observe variants at target probabilities across allele frequencies.

### SV annotation and gene-overlap classification

SV coordinates were intersected with gene annotations from the Ssal_v3.1 gene annotation file using BEDTools (Quinlan and Hall 2010). Intersections were used to classify SVs according to their relationship with annotated gene features, including gene body, exon, coding sequence, untranslated region, and nearby or intergenic positions where applicable.

For SV-eQTL candidates, gene-overlap categories were defined relative to the associated target gene. Cis SV-eQTLs were classified as target-gene-overlapping when the SV intersected the associated gene body, as other-gene-overlapping when the SV intersected an annotated gene other than the associated target gene, and as no gene-body overlap when the SV did not intersect annotated gene bodies. Trans SV-eQTLs were similarly classified according to whether they overlapped any annotated gene body.

### SV-eQTL mapping

SV-eQTL mapping was performed using QTLtools v1.3.1 (Delaneau et al. 2017) with imputed SV dosages and normalized gill gene expression from the 906 offspring. The covariate matrix included father, mother, sex, tank, weight, body length, condition factor, skin colouration, and ten genomic principal components. These covariates were included to account for family structure, experimental design, phenotypic covariates, and genome-wide ancestry.

Cis-eQTL mapping was performed using a permutation-based approach with 5000 permutations. Associations were classified as cis when the SV was located within 5 Mb of the tested gene. Trans-eQTL mapping was performed using an approximation approach in which 1000 phenotypes were randomly sampled to estimate the null distribution for multiple-testing correction. Associations were classified as trans when the SV was located more than 5 Mb from the tested gene or on a different chromosome. SV-eQTL candidates were retained after multiple-testing correction and filtering for imputation quality.

### SNP-eQTL comparison and conditional regression

SV-eQTL candidates were compared with SNP-eQTL results generated previously from the same gill RNA-seq cohort. SNP-eQTL lead SNPs were identified for corresponding target genes where available. SV-eQTL candidates without an available or non-monomorphic SNP-eQTL lead SNP were excluded from the lead-SNP conditional analysis.

Local SNP tagging of SVs was evaluated by calculating linkage disequilibrium between each SV and nearby SNPs. Mean LD was summarized within ±10 kb, ±100 kb, and ±1 Mb windows. For each SV, the nearby SNP with the highest LD within ±1 Mb was also identified. SV-eQTL candidates were compared with non-eQTL SVs using Wilcoxon rank-sum tests, with Benjamini-Hochberg correction across distance windows.

Conditional regression was used to evaluate whether SV-eQTL signals were retained after accounting for nearby SNP signals. For each SV-eQTL candidate, gene expression was modelled as a function of both SV dosage and SNP genotype or dosage in the same regression model:

Expressionᵢ = β₀ + βSV SV dosageᵢ + βSNP SNP dosageᵢ + covariatesᵢ + εᵢ

The purpose of this conditional regression analysis was to assess haplotype representation rather than to select the best-fitting predictive model. Specifically, we asked whether the imputed SV dosage and nearby SNP markers captured the same expression-associated haplotype signal, and whether the SV association remained after accounting for individual SNP markers.

Two SNP definitions were used. First, where available, the SNP-eQTL lead SNP for the corresponding target gene was included as the conditioning SNP. Second, the nearby SNP with the highest LD to the SV within ±1 Mb was used as a local tagging SNP. The first analysis tested whether SV signals were retained after accounting for the strongest SNP-eQTL marker previously detected for the target gene. The second analysis tested whether SV signals were retained after accounting for the best available local SNP tag.

SV and SNP terms were considered retained at a Benjamini-Hochberg adjusted P value threshold of 0.05. Conditional outcomes were classified as only SV retained, both SV and SNP retained, only SNP retained, or neither retained. Paired Wilcoxon tests were used to compare conditional P values and absolute effect sizes between SV and SNP terms. McNemar’s test was used to compare the frequency of SV-only and SNP-only retention.

### Representative configurations and statistical analyses

Representative SV-eQTL candidates were selected to illustrate distinct genomic configurations, including direct target-gene overlap, nearby cis association without target-gene overlap, trans association, short-read-detected insertion, and positive or negative expression effects. Because the offspring cohort had SNP-array genotypes and RNA-seq reads but not WGS, RNA-seq alignments were used to visualize transcript context at selected loci.

All statistical analyses and visualizations were performed in R version 4.6.0. Multiple testing was controlled using the Benjamini-Hochberg procedure. Genome interval analyses used GenomicRanges and BEDTools. Plots were generated using ggplot2 and assembled using patchwork. Additional R packages included readr, dplyr, tidyr, stringr, rtracklayer, scales, and forcats.

#### Statistical analysis and visualization

All statistical analyses and visualizations were performed in R. Multiple testing was controlled using the Benjamini-Hochberg procedure unless otherwise stated. Genome interval operations were performed using BEDTools and R packages for genomic ranges. Plots were generated using ggplot2 and assembled using patchwork. Additional R packages included readr, dplyr, tidyr, stringr, rtracklayer, scales, and forcats.

## Results

### The imputed SV catalogue captures genome-wide structural variation in Atlantic salmon

The combined SV catalogue contained 114,293 variants before size filtering (**Figure 1**, **Figure 2)**. The catalogue was dominated by long-read-derived variants, and deletions represented the largest class overall. When variants larger than 50 bp were considered, the SV catalogue contained 100,269 variants. Of these, 97,942 variants (97.7%) were derived from long-read-based discovery and 2,327 variants (2.3%) from a stringent short-read-based discovery (Diblasi et al. 2026). Deletions represented the largest SV class, accounting for 97,790 of 97,942 long-read-derived variants (99.8%) and 1,458 of 2,327 short-read-derived variants (62.7%). Inversions were detected only among the long-read-derived SVs and were rare (31 variants). Among short-read-derived SVs, deletions and insertions were the main classes, with 1,458 deletions (62.7%) and 869 insertions (37.3%) detected. However, long-read-derived duplications showed a distinct size distribution: 11,892 of 12,013 long-read-derived duplications (99.0%) were ≤50 bp, whereas only 121 (1.0%) were larger than 50 bp. We therefore separated variants by size and focused subsequent SV summaries on variants larger than 50 bp (**Figure 2**).

**Figure 2.**
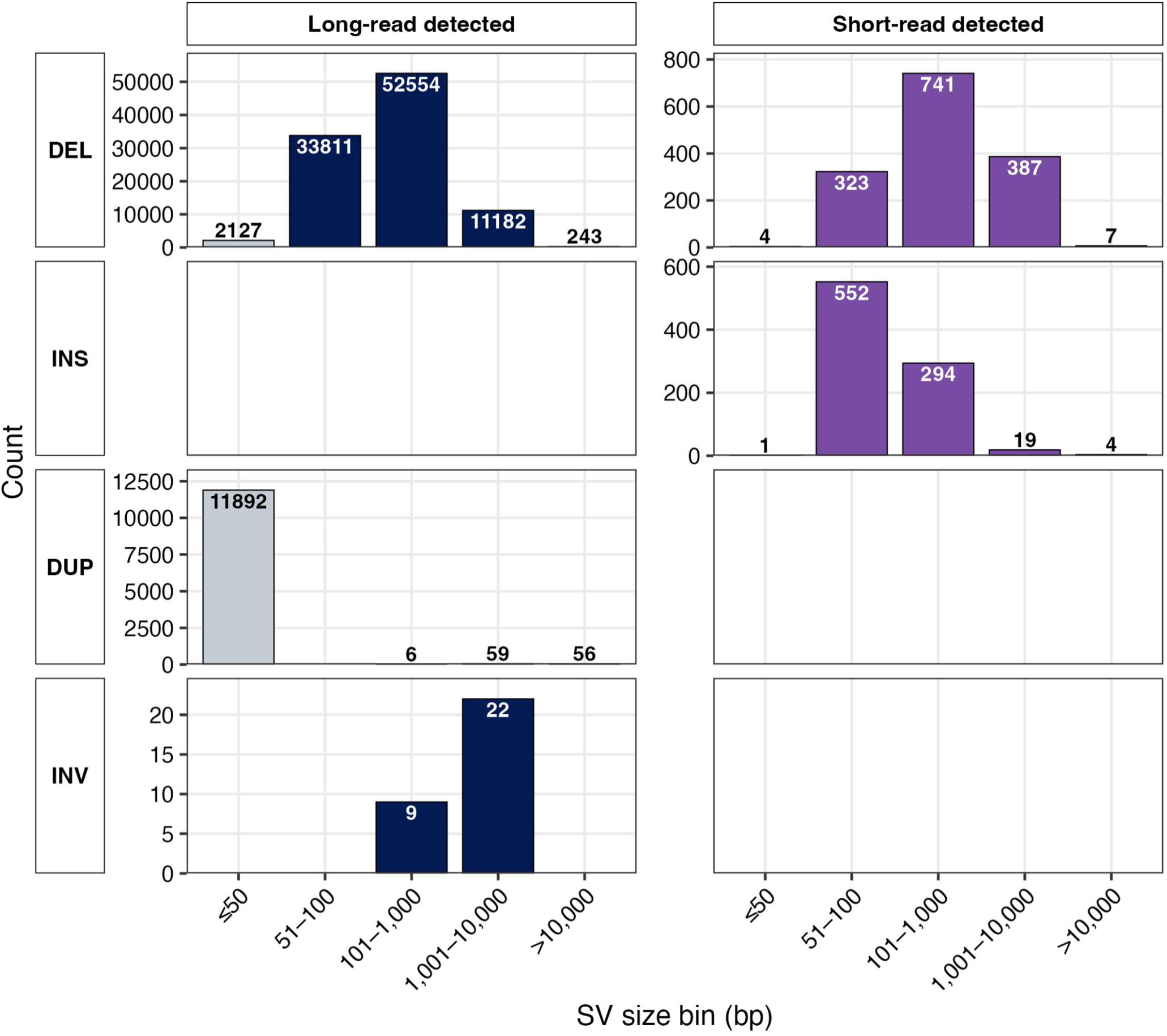
Size distribution of structural variants by variant class and detection source. Structural variants were grouped by size bin and variant type, and counts were compared between long-read-detected and short-read-detected variants. Bars show the number of variants in each size class for deletions (DEL), insertions (INS), duplications (DUP), and inversions (INV). Variants of ≤50 bp are shown in grey, while larger long-read-detected and short-read-detected variants are shown in navy and purple, respectively. Counts are indicated above or within bars. Y-axis scales are shown independently for each panel to accommodate differences in variant abundance across classes.

We also compared the imputed SV dataset with independently identified inversion-like regions detected by a SNP-based sliding-window PCA approach. Using direct coordinate overlap, 778 overlaps were detected between imputed SVs and 51 inversion-like regions (**Table S1**) (Diblasi et al. 2026). These overlaps involved 771 unique imputed SVs, most of which were deletions (700 DEL, 54 DUP, 14 INS, and 3 INV). Thus, the independently identified inversion-like regions contained multiple imputed SVs, mainly deletions.

Atlantic salmon has a strongly sex-specific recombination landscape, with male recombination being spatially restricted to the telomeric ends of the chromosomes, and female recombination being relatively less restricted but elevated towards the centromeres (Sardell and Kirkpatrick 2020; Brekke et al. 2023; MacLeod-Bigley and Boulding 2023). Because recombination rate can shape the accumulation, persistence, and genomic distribution of structural variation, we examined whether SV density varied with local recombination rate in a sex-specific manner. To do this, we summarized SV counts in 1 Mb genomic windows and compared SV density across quartiles of male and female recombination rate (**Table S2)**.

SV density differed significantly among sex-specific recombination-rate quartiles (**Figure 3A**) (Kruskal-Wallis test, female: P = 8.26 × 10⁻⁴¹; male: P = 2.78 × 10⁻⁷⁵). In the female map, windows in the lowest recombination-rate quartile showed the highest SV density, with a median of 65 SVs per 1 Mb window, compared with 24 to 27 SVs in the other female quartiles. In contrast, in the male map, windows in the highest recombination-rate quartile showed the highest SV density, with a median of 70 SVs per 1 Mb window, compared with 24 to 26 SVs in the other male quartiles. Thus, SV density was associated with sex-specific recombination landscapes, with elevated SV density in regions of low female recombination and high male recombination.

**Figure 3.**
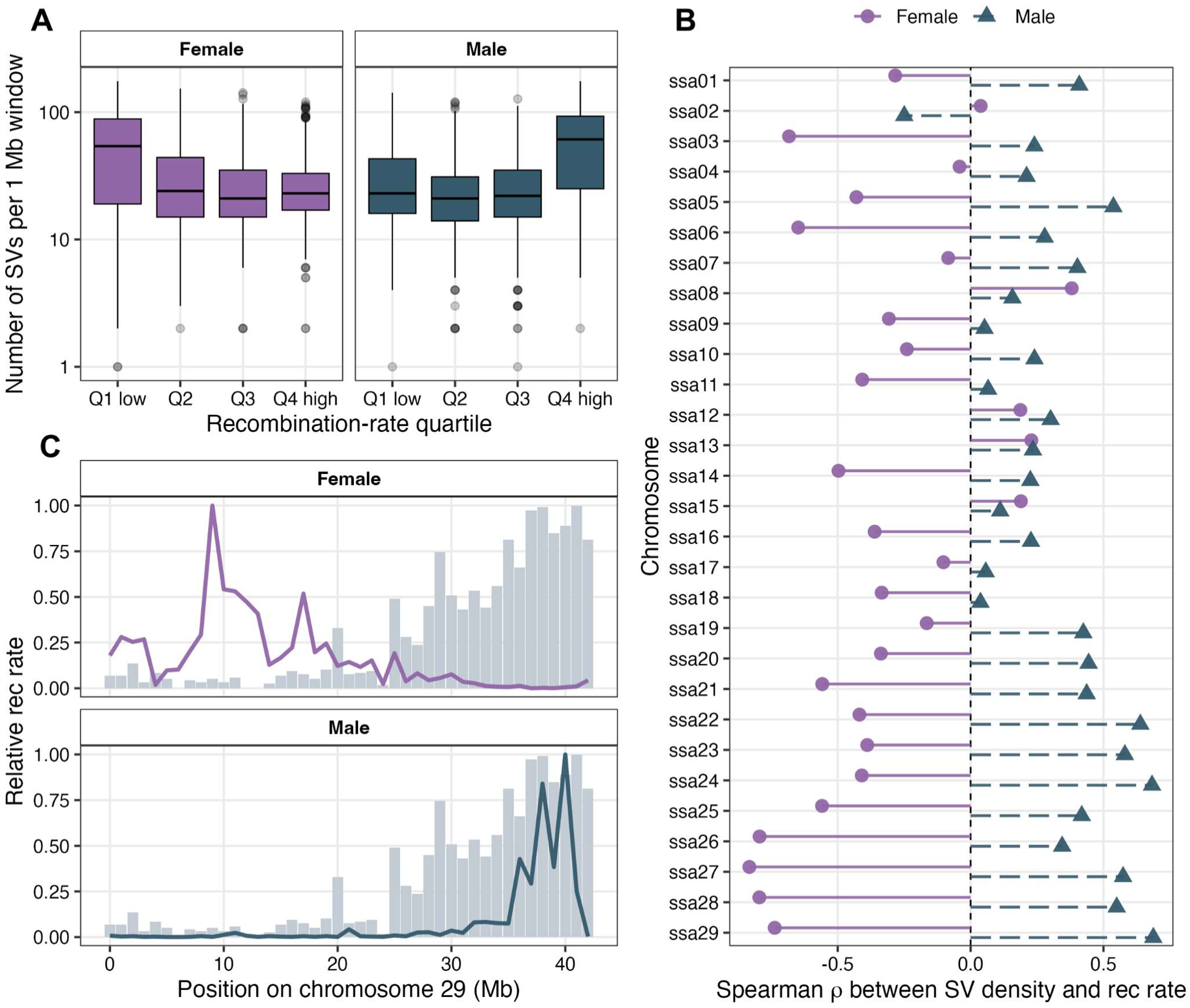
Relationship between structural variant density and sex-specific recombination rate across the Atlantic salmon genome. **A**, Distribution of the number of SVs >50 bp per 1 Mb window across sex-specific recombination-rate quartiles. Windows were grouped separately for female and male recombination rates. Boxes show the interquartile range, horizontal lines indicate medians, whiskers show the non-outlier range, and points indicate outliers. **B**, Chromosome-wise Spearman correlations between SV density and recombination rate. Female and male recombination-rate correlations are shown separately for each chromosome. The dashed vertical line indicates ρ = 0. **C**, Local distribution of SV density and recombination rate along chromosome 29. Grey bars show scaled SV density per 1 Mb window, and lines show scaled female or male recombination rate within the chromosome. Values were scaled within the chromosome to compare local patterns between SV density and recombination rate.

To examine chromosome-specific patterns, we calculated Spearman correlations between SV density and recombination rate separately for each chromosome and sex (**Figure 3B**). Seventeen chromosomes showed significant opposite associations between the female and male recombination maps, with negative correlations in females and positive correlations in males. Chromosome 29 showed the strongest male–female contrast, with SV density negatively correlated with female recombination rate (Spearman’s ρ = −0.730, P = 2.79 × 10⁻⁸) but positively correlated with male recombination rate (ρ = 0.700, P = 1.75 × 10⁻⁷) (**Figure 3 C**).

### SV-eQTL candidates are enriched for short-read-derived variants

We next asked whether a subset of these SVs also has detectable regulatory effects on gene expression. To address this, we performed SV-based eQTL mapping using imputed SV genotypes and transcriptomic data. We applied QTLTools-based cis and trans eQTL mapping across 906 individuals using imputed SV genotypes and transcriptomic data. After correction for multiple-testing bias and filtering on imputation quality (average imputation R² > 0.5), 51 unique SVs were significantly associated with expression variation (**Figure 4A, Table S3**). Gene-overlapping variants were modestly enriched among SV-eQTL candidates: 39 of 51 SV-eQTL candidates overlapped annotated genes, compared with 71,343 of 114,293 variants in the full imputed set (Fisher’s exact test, odds ratio = 1.96, p = 0.042).

**Figure 4.**
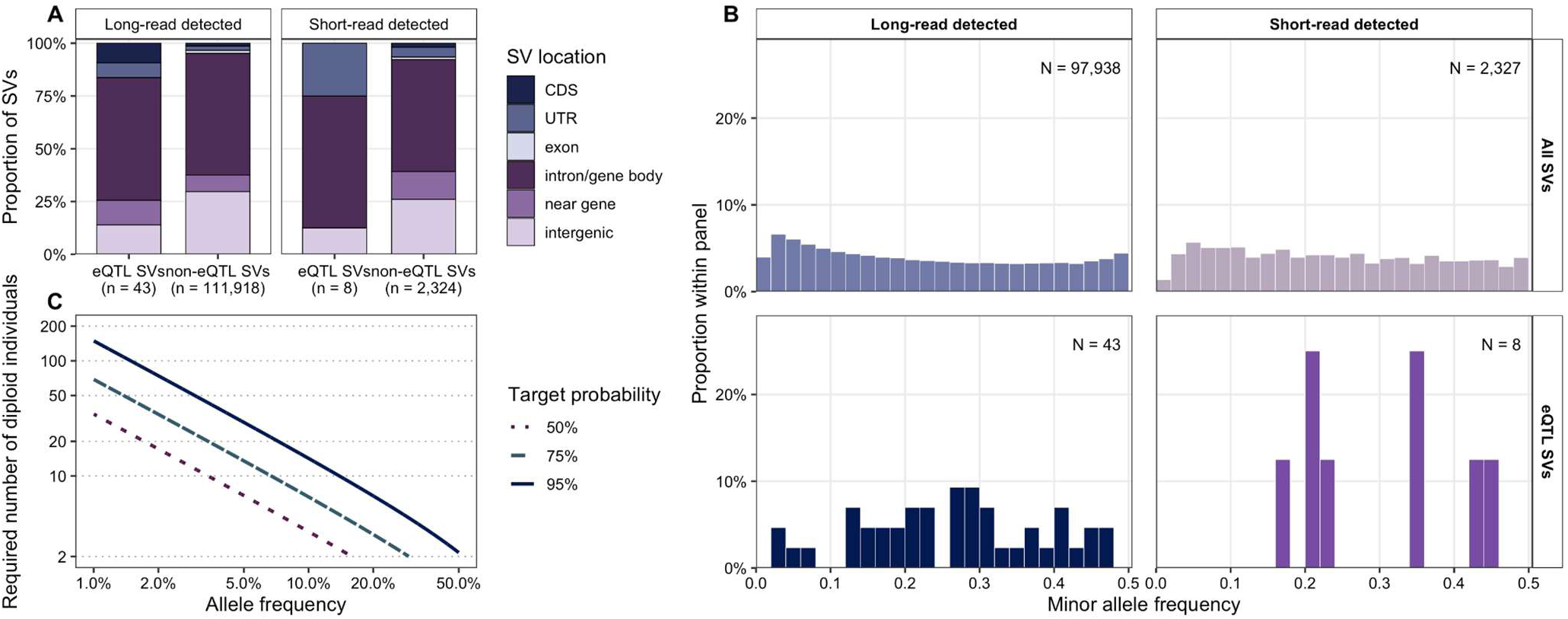
Genomic distribution, allele-frequency spectrum, and discovery-panel requirements of structural variants. **A** Proportional distribution of SV genomic locations for eQTL SVs and non-eQTL SVs, shown separately for long-read-detected and short-read-detected variants. SVs were classified as CDS, UTR, exon, intron/gene body, near gene, or intergenic. Numbers in parentheses indicate the total number of SVs in each group. **B** Minor allele frequency distributions of all SVs and eQTL SVs, shown separately for long-read-detected and short-read-detected variants. Bars indicate the proportion of variants within each panel, and panel labels indicate the number of variants with allele-frequency estimates. **C** Expected number of diploid individuals required in a discovery panel to observe at least one alternate allele at target probabilities of 50%, 75%, and 95%, shown as a function of allele frequency. Curves illustrate the rapid increase in required sample size as allele frequency decreases.

Short-read-detected SVs were enriched among SV-eQTL candidates. Although they represented only 2.0% of the full imputed variant set (2,332/114,293), they accounted for 15.7% of SV-eQTL candidates (8/51; Fisher’s exact test, odds ratio = 8.96, p = 8.66 × 10⁻⁶). Thus, supplementing the long-read catalog with short-read-detected variants recovered expression-associated SVs that would have been missed by a long-read-only catalog.

To assess whether this enrichment could reflect allele-frequency ascertainment during variant discovery, we compared the minor allele frequency distributions of long-read- and short-read-detected SVs and evaluated the expected probability of observing variants as a function of discovery panel size (**Figure 4B**, **Figure 4C**). The MAF distributions differed between long-read- and short-read-detected SVs (Wilcoxon rank-sum test, P = 6.61 × 10⁻¹⁴; Kolmogorov–Smirnov test, D = 0.084, P = 9.12 × 10⁻¹⁵). Short-read-detected SVs were not enriched for rare alleles; their median MAF was slightly higher than that of long-read-detected SVs (0.228 vs. 0.211), and the proportion of rare variants was lower among short-read-detected SVs both at MAF < 0.01 (0.34% vs. 1.43%) and MAF < 0.05 (8.72% vs. 13.63%). Conversely, common variants were slightly more frequent among short-read-detected SVs, with 78.47% having MAF ≥ 0.10 compared with 72.94% of long-read-detected SVs.

Among SV-eQTL candidates, the short-read-detected subset was small (n = 8), but showed a similar descriptive pattern, with a median MAF of 0.283 compared with 0.273 for long-read-detected SV-eQTL candidates. Thus, the enrichment of short-read-detected SVs among SV-eQTL candidates was not explained by an excess of rare alleles among the short-read-detected variants retained after filtering and imputation (**Figure 4B**).

This pattern is consistent with the expected ascertainment imposed by small discovery panels. Under a simple diploid sampling model (N = 2 for long-read sequencing, and N = 112 for short- read sequencing), the probability of observing at least one alternate allele is P = 1 - (1 - p)^(2N), where p is allele frequency and N is the number of individuals in the discovery panel. This model showed that a two-individual discovery panel has limited probability of capturing variants unless they are common. For example, observing at least one alternate allele with 95% probability would require approximately 150 individuals for a variant at 1% allele frequency, 75 individuals at 2%, 30 individuals at 5%, and 15 individuals at 10% (**Figure 4C**). Thus, a two-individual discovery panel would be expected to capture only relatively common variants, whereas a discovery panel of several hundred individuals would be sufficient to capture variants segregating at low to moderate frequencies with high probability.

### SV-eQTL candidates occupy diverse genomic contexts and are only partially represented by nearby SNPs

We next examined the genomic context of the 51 SV-eQTL candidates to assess the possible routes by which these variants were associated with gene expression. This analysis was not limited to whether an SV directly overlapped its associated target gene, but also considered overlap with other annotated gene bodies, the absence of gene-body overlap, cis versus trans classification, and the direction of the expression effect. Among the 35 cis-eQTL SVs, only 9 overlapped the target gene body, whereas 19 overlapped another annotated gene and 7 did not overlap any gene body (**Figure 5A**). All target-gene-overlapping cis-eQTL SVs were deletions, but their effects were not uniformly negative: 4 had negative slopes and 5 had positive slopes (**Figure 5B**). Among trans-eQTL SVs, 11 of 16 overlapped another annotated gene, whereas 5 showed no gene-body overlap. These observations indicate that the 51 SV-eQTL candidates did not represent a single simple class, such as “deletions directly overlapping their target genes and reducing expression”. Instead, the candidates included target-gene-overlapping cis deletions, cis variants located in other gene bodies, cis variants outside annotated gene bodies, and trans variants with or without overlap to other annotated genes.

**Figure 5.**
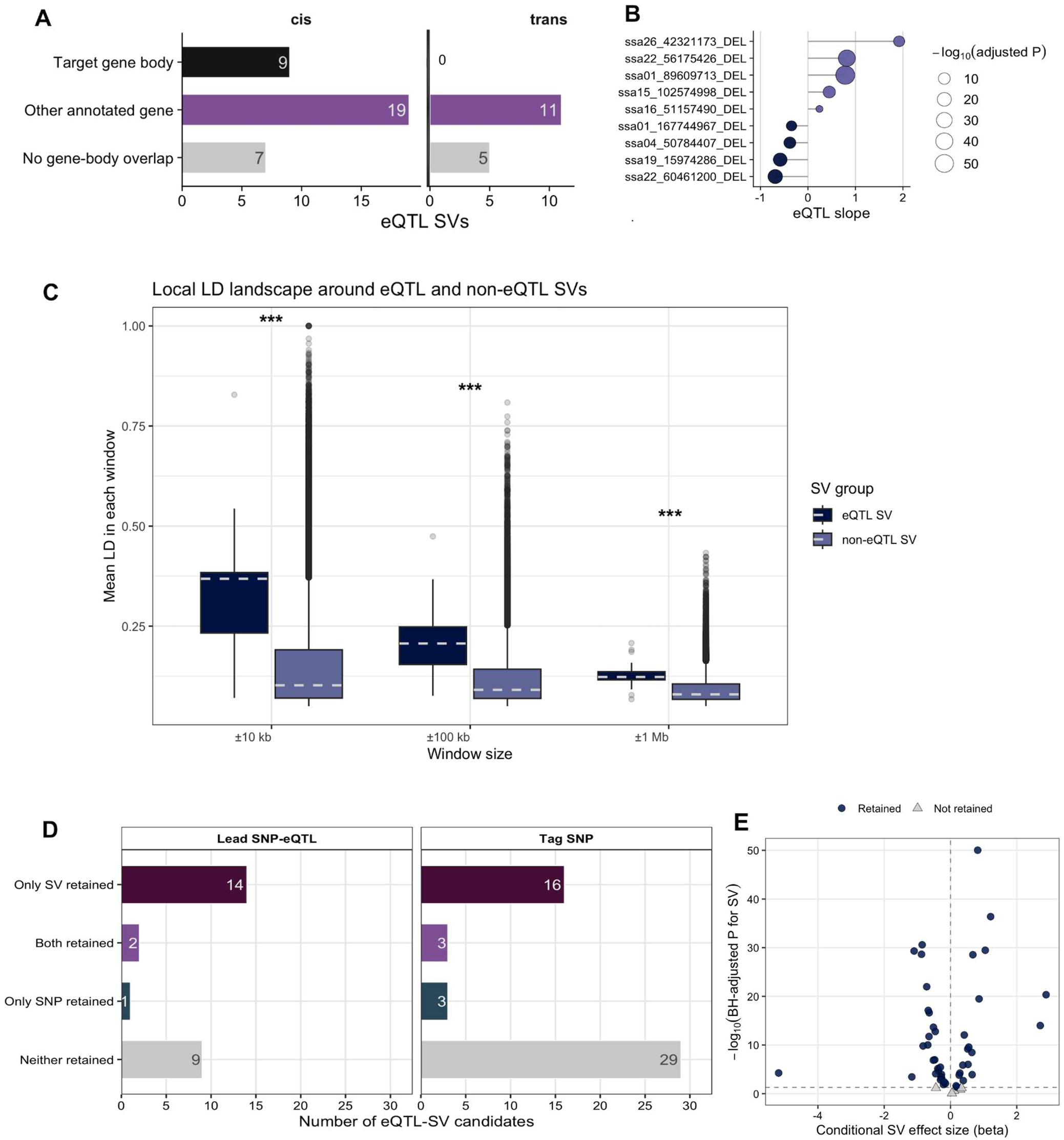
Structural variants associated with gene expression, local LD patterns, and conditional independence from nearby SNPs. (A) Gene-overlap classification of SV-eQTL candidates, separated into cis and trans associations. Bars show the number of SVs overlapping the target gene body, overlapping another annotated gene, or showing no gene-body overlap. (B) Effect sizes of gene-overlapping deletion-eQTLs. Points show the estimated eQTL slope for each SV, and point size indicates −log10(BH-adjusted P value). (C) Local linkage disequilibrium around eQTL SVs and non-eQTL SVs. Mean LD within ±10 kb, ±100 kb, and ±1 Mb windows was compared between eQTL SVs and non-eQTL SVs. Boxplots show the distribution across SVs, with dashed horizontal lines indicating group means. Asterisks indicate significant differences between groups. (D) Conditional regression outcomes after accounting for nearby SNPs. SV associations were tested after conditioning on either the lead SNP-eQTL or the highest-LD nearby tagging SNP. Bars show the number of SV-eQTL candidates for which only the SV was retained, both the SV and SNP were retained, only the SNP was retained, or neither was retained. (E) Conditional SV effects after conditioning on the highest-LD nearby SNP. Points show the conditional SV regression coefficient and −log10 BH-adjusted P value for the SV term. Retained associations are defined as SV terms with BH-adjusted P < 0.05 and are shown as circles; non-retained associations are shown as triangles. The horizontal dashed line indicates the BH-adjusted significance threshold of 0.05, and the vertical dashed line indicates zero conditional SV effect.

Because these genomic configurations alone do not establish whether the observed expression associations are attributable to the SVs themselves or to linked SNPs on the same local haplotypes, we next evaluated local SNP tagging and conditional association patterns. Structural variants are often correlated with nearby SNPs because they share local haplotypes. If an SV-eQTL signal is well tagged by nearby SNPs, the same regulatory association may already be captured by standard SNP-based eQTL analysis. In that case, adding SV genotypes would provide limited additional information. Conversely, if the SV is only partially tagged by SNPs, or if the SV signal remains after accounting for nearby SNPs, this would indicate that SV genotyping captures regulatory variation that is not fully represented by SNP markers. We therefore asked two related questions. First, are SV-eQTL candidates more strongly tagged by nearby SNPs than other SVs? Second, when both the SV and a nearby SNP are included in the same model, which signal is retained?

To evaluate whether SV-eQTL candidates were likely to be captured by nearby SNP markers, we first compared local SNP tagging between SV-eQTL candidates and non-eQTL SVs. If SV-eQTL candidates were generally well tagged by nearby SNPs, their expression associations could be largely recoverable from SNP-based eQTL analysis. Conversely, incomplete local tagging would indicate that SV-based genotyping captures regulatory variation that is only partially represented by nearby SNPs.

We compared SV-eQTL candidates with non-eQTL SVs using the subset of variants for which local SNP LD information was available. Local LD could be calculated for 31 of the 51 SV-eQTL candidates and for 49,873 non-eQTL SVs. SV-eQTL candidates showed higher average LD with nearby SNPs than non-eQTL SVs across all three distance windows (**Figure 5C**). The median mean LD for SV-eQTL candidates was 0.368 within 10 kb, 0.207 within 100 kb, and 0.123 within 1 Mb, compared with 0.102, 0.091, and 0.080, respectively, for non-eQTL SVs. These differences were significant for all three windows (Wilcoxon rank-sum test, BH-adjusted p = 4.36 × 10⁻¹⁰ for 10 kb, 4.24 × 10⁻¹⁰ for 100 kb, and 7.69 × 10⁻¹⁰ for 1 Mb). Although SV-eQTL candidates tended to occur in regions with higher local SNP tagging than non-eQTL SVs, the average LD values remained moderate, particularly over the 1 Mb window. The corresponding boxplot shows consistently higher local SNP LD for SV-eQTL candidates than for non-eQTL SVs across the 10 kb, 100 kb, and 1 Mb windows.

### Conditional regression shows that individual SNP markers do not consistently capture imputed SV-eQTL signals

Higher local LD alone does not show whether the expression association is driven by the SV or by a linked SNP. We therefore performed conditional regression analyses in which the SV genotype and a nearby SNP genotype were included in the same model (**Table S4**). This analysis tested whether the SV still explained expression variation after accounting for the SNP, and conversely whether the SNP still explained expression variation after accounting for the SV. We used two SNP definitions: the lead SNP from the SNP-eQTL analysis where available, and the nearby SNP showing the highest LD with each SV within ±1 Mb. The first analysis tested whether the SV signal remained after conditioning on the strongest SNP-eQTL signal previously detected at the locus (Diblasi et al. 2025), whereas the second provided a more conservative test based on the best available local SNP tag.

For the SNP-eQTL lead analysis, 26 SV-eQTL candidates had an available and non-monomorphic SNP-eQTL lead SNP. Using a BH-adjusted threshold of 0.05, the SV term was retained in 16 of 26 candidates (61.5%), whereas the SNP term was retained in 1 of 26 candidates (3.8%). Fifteen candidates retained only the SV term, one candidate retained both terms, and ten candidates retained neither term. No candidate retained only the SNP term (**Figure 5D**). The asymmetry between SV-only and SNP-only retention was significant by McNemar’s test (p = 0.00195). The SV terms were also stronger than the SNP terms when analysed as paired tests. In the SNP-eQTL lead analysis, the median conditional p-value was 0.00348 for the SV term and 0.292 for the SNP term. A paired Wilcoxon test comparing SV and SNP conditional p-values was significant (p = 1.66 × 10⁻⁴). Conditional effect sizes showed the same pattern: the median absolute effect size was 0.419 for SVs and 0.120 for SNPs, with a paired Wilcoxon p-value of 3.28 × 10⁻⁶. Thus, among testable loci with an available SNP-eQTL lead SNP, SV effects were more often retained and generally stronger than the corresponding SNP effects.

As a complementary analysis, we repeated the conditional regressions using the nearby SNP with the highest LD to each SV. This analysis could be performed for all 51 SV-eQTL candidates. Using the same BH-adjusted threshold, the SV term was retained in 14 of 51 candidates (27.5%), whereas no SNP term was retained. Fourteen candidates retained only the SV term, and 37 candidates retained neither term (**Figure 5D**, **Figure 5E**). The SV-only category was again more frequent than the SNP-only category (McNemar’s test, p = 0.00591). The median conditional p-value for the SV term increased from 0.00348 in the SNP-eQTL lead analysis to 0.111 in the max-LD SNP analysis, consistent with stronger local SNP tagging. Nevertheless, SV terms still showed lower conditional p-values than SNP terms in paired comparison (median p = 0.111 for SVs vs. 0.326 for SNPs; paired Wilcoxon p = 0.0454). Median absolute effect sizes were also larger for SVs than for max-LD SNPs (0.520 vs. 0.411; paired Wilcoxon p = 0.00880).

Together, these conditional analyses indicate that a subset of SV-eQTL signals persisted after SNP conditioning, particularly when conditioning on SNP-eQTL lead SNPs, whereas conditioning on the strongest local tagging SNP attenuated many SV associations. These retained associations were then examined as representative examples spanning distinct genomic configurations, including direct target-gene overlap, nearby cis effects without target-gene overlap, trans associations, and short-read-detected insertions.

### SV-eQTLs as possible multiple configurations of regulatory haplotypes

Together, the conditional analyses indicate that a subset of SV-eQTL signals persisted after SNP conditioning, particularly when conditioning on SNP-eQTL lead SNPs, whereas conditioning on the strongest local tagging SNP attenuated many SV associations. We therefore examined retained SV-eQTL candidates as representative examples of the structural configurations through which imputed SV alleles were associated with gene expression (**Figure 6**). These examples were not interpreted as definitive evidence that the SV itself was causal. Rather, they were used to place retained SV-eQTL signals into observed genomic configurations, including direct target-gene-overlapping deletions, nearby cis deletions without target-gene overlap, distal trans deletions, and short-read-detected cis insertions with opposite directions of expression effect.

**Figure 6.**
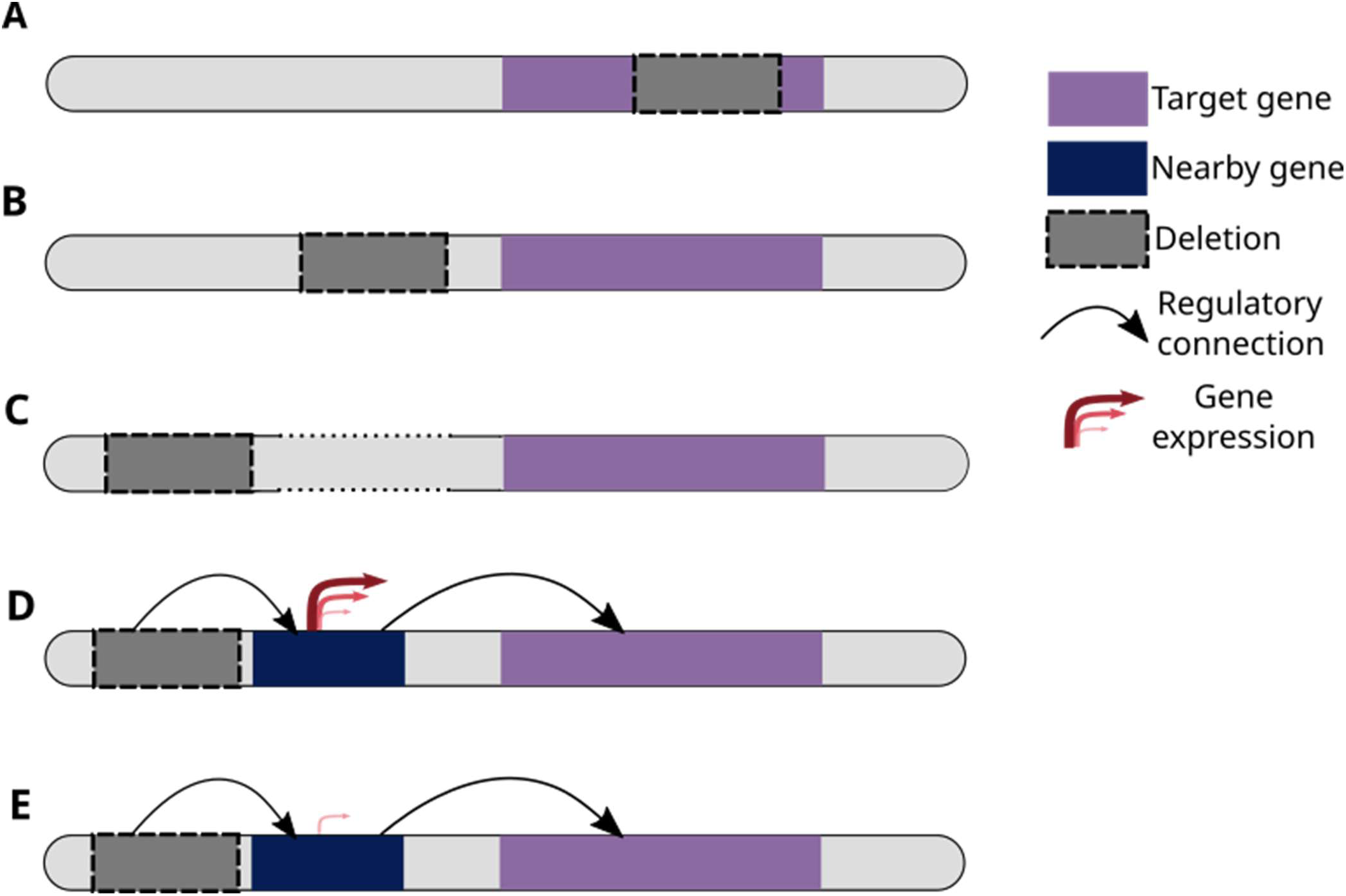
Schematic summary of observed configurations among retained SV-eQTL associations. **(A)** Direct target-gene-overlapping deletion. The deletion intersects the annotated body of the associated gene, representing a configuration consistent with altered transcript structure, untranslated-region sequence, gene dosage, or nearby regulatory sequence. **(B)** Nearby cis deletion without target-gene overlap. The deletion lies close to the associated target gene but outside its annotated gene body. In this configuration, the deleted interval can be intergenic or can overlap another annotated gene, indicating a local regulatory haplotype rather than direct disruption of the target gene itself. **(C)** Distal trans deletion. The deletion is located far from the associated target gene and does not define a simple local overlap model, representing a configuration consistent with distal or indirect regulatory effects. **(D)** Short-read-detected cis insertion with positive expression effect. The insertion is associated with increased expression of a nearby target gene, representing a small structural variant within a local regulatory haplotype. **(E)** Short-read-detected cis insertion with negative expression effect. The insertion is associated with decreased expression of a nearby target gene, indicating that short-read-detected structural variants can be linked to either direction of expression effect.

For selected representative loci, we also inspected RNA-seq alignments from the offspring cohort to provide transcript context for the SV-eQTL signals. The offspring had SNP-array genotypes and RNA-seq data, but not whole-genome sequencing; therefore, these RNA-seq views were not used to validate SV breakpoints or SV genotypes. Instead, they were used to examine, where visible from RNA-seq reads, transcript coverage, splice-junction patterns, and the position of eQTL-associated SV intervals relative to annotated gene models (**Supplementary Figure 1**).

One direct cis example was a 1,936 bp deletion on chromosome 1 (ssa01:89609713:DG) associated with ENSSSAG00000102775, a gene likely corresponding to PGRMC1 (**Figure 6A**). This association had complete imputation accuracy (R² = 1.00) and showed a strong genotype-associated expression difference (Wilcoxon FDR = 8.71 × 10⁻¹⁷). The variant overlapped the target gene body, and VEP annotated consequences including feature truncation and 3′ UTR-related effects. This locus therefore represents a direct target-gene-overlapping deletion, in which the imputed structural variant intersects the associated gene.

A second cis example showed a different local configuration. A 4,431 bp deletion on chromosome 24 (ssa24:47377244:DG) was associated with ENSSSAG00000063251, annotated as pias2 (**Figure 6B**). This SV had acceptable imputation accuracy (R² = 0.687) and was associated with a significant expression difference (Wilcoxon FDR = 1.12 × 10⁻¹³). In contrast to the PGRMC1-like example, VEP annotated this variant as downstream of the target gene rather than as a direct target-gene overlap. This locus therefore represents a nearby cis deletion without target-gene overlap. In this configuration, the SV may be intergenic relative to the target gene or may overlap another annotated feature, but the expression association is not explained by direct disruption of the target gene body.

The candidate set also included distal trans configurations. A 4,732 bp deletion on chromosome 10 (ssa10:43236855:DG) was identified as a trans-eQTL for ENSSSAG00000067074 (**Figure 6C**). This association had high imputation accuracy (R² = 0.808) and a significant expression difference (Wilcoxon FDR = 2.00 × 10⁻²⁹). The target gene showed similarity to E-selectin-related genes, providing an example of a distal SV-eQTL association involving a gene with a possible immune or inflammatory annotation. Because the SV and target gene are not in a simple local overlap configuration, this class was treated as a distal or indirect regulatory association.

Short-read-detected insertions provided additional configurations not represented by the long-read-derived deletion examples. One example was ssa11_102108843_SR, an 85 bp insertion on chromosome 11. This was identified as a cis-eQTL for ENSSSAG00000007780, annotated as sparta (**Figure 6D**). The association had complete imputation accuracy (R² = 1.00), a significant genotype-associated expression difference (Wilcoxon FDR = 3.38 × 10⁻¹²), and a positive eQTL slope (slope = 0.445), indicating higher expression associated with the alternate/SV allele. The insertion did not overlap the sparta gene body. Instead, it overlapped another annotated gene body and was located approximately 698 kb upstream of the sparta coordinates. This locus therefore represents a short-read-detected cis insertion associated with increased target-gene expression.

A second short-read-detected cis insertion showed the opposite effect direction. ssa15_33256881_SR, a 51 bp insertion on chromosome 15, was associated with ENSSSAG00000040719, annotated as slc35a1 (**Figure 6E**). This association had high imputation accuracy (R² = 0.939), a significant expression difference (Wilcoxon FDR = 2.12 × 10⁻⁸), and a negative eQTL slope (slope = −0.423), indicating lower expression associated with the alternate/SV allele. As in the sparta example, the insertion did not overlap the target gene body but overlapped another annotated gene body. It was located approximately 19.9 kb downstream of the slc35a1 coordinates. Together, the sparta and slc35a1 examples show that short-read-detected cis insertions were associated with both increased and decreased expression, indicating that SV-eQTLs cannot be reduced to a single loss-of-expression model.

Finally, ssa07_19430663_SR, a 62 bp short-read-detected insertion on chromosome 7, illustrated a short-read-derived trans association without annotated gene-body overlap. This SV was associated with ENSSSAG00000091100, a protein-coding gene without a gene name in the current annotation table. The association had high imputation accuracy (R² = 0.955), a significant expression difference (Wilcoxon FDR = 1.66 × 10⁻⁶), and a negative eQTL slope (slope = −0.194). The insertion did not overlap any annotated gene body and was located approximately 15.03 Mb from the target gene coordinates, consistent with its classification as a trans-eQTL. This example was not included as a separate configuration in **Figure 6**, but it further supports the observation that retained SV-eQTLs included short-read-derived and distal associations beyond direct target-gene overlap.

Overall, these representative loci show that retained SV-eQTLs occurred across multiple observed configurations rather than being restricted to direct disruption of the target gene. Deletion examples included direct target-gene overlap, nearby cis association without target-gene overlap, and distal trans association. Short-read-detected insertion examples further showed both positive and negative expression effects. These patterns support the interpretation that retained SV-eQTLs represent structurally defined components of regulatory haplotypes rather than a homogeneous class of disruptive variants.

## Discussion

Structural variants are increasingly recognized as important contributors to gene expression variation, with previous studies showing that deletions, insertions, duplications, inversions, and repeat-associated variants can affect transcript abundance, gene regulation, and complex trait-associated regulatory architecture (Chiang et al. 2017; Scott et al. 2021). However, the interpretive problem addressed here is how graph-genotyped SVs should be interpreted when they are imputed into a SNP-genotyped expression cohort. In human and other well-resourced systems, SV-eQTL studies have often linked directly genotyped or whole-genome-sequenced SVs to gene expression, allowing SV-expression associations to be evaluated in the same individuals in which SV genotypes were observed (Noyvert et al. 2025; Mapel et al. 2026). By contrast, many large expression cohorts, especially outside human genetics, are still genotyped primarily with SNP arrays or short-read-derived SNP panels, whereas comprehensive SV discovery and direct SV genotyping remain limited to smaller reference sets. This creates a common gap between the scale of available transcriptomic phenotypes and the scale at which structural variants can be directly observed. Here, we addressed this gap by graph-genotyping SVs in a WGS reference panel and projecting these structurally defined alleles into a large RNA-seq cohort through haplotype-based imputation. Thus, the main contribution of this study is not simply adding SVs as another variant class, but using imputed SV dosages to assign structural interpretation to regulatory haplotypes that would otherwise be represented only by SNP markers.

In this study, SV genotypes in the expression cohort were not directly observed but inferred by haplotype-based imputation from a WGS reference panel. Studies in humans and other well-resourced systems have linked directly genotyped or whole-genome-sequenced SVs to gene expression, often emphasizing the functional consequences of variants that intersect genes, regulatory elements, or chromatin contacts. Those studies establish that SVs can be important regulatory variants. In contrast, our design addresses a different and common setting: a large RNA-seq cohort with SNP-array genotypes, but without whole-genome sequencing for the same individuals. The resulting SV-eQTL signals should therefore not be interpreted as direct evidence that the imputed SV allele is the causal regulatory variant. Rather, they should be interpreted as regulatory haplotypes whose support depends on imputation accuracy, local haplotype structure, allele frequency, and the extent to which nearby SNP markers capture the same signal. This haplotype-based interpretation is consistent with recent work arguing that regulatory gene-expression associations can reflect haplotype effects rather than single causal-variant effects (Greenwood et al. 2025). It is also consistent with known limitations of eQTL fine-mapping and conditional analysis when linked or multiple local regulatory variants are present (Zeng et al. 2017; Kundu et al. 2022; Li and Zhou 2025). Thus, imputed SVs can make SNP-defined eQTL haplotypes more biologically interpretable, but they do not by themselves prove that the SV allele is causal.

Consistent with the haplotype-based nature of SV imputation, SV-eQTL candidates were more strongly tagged by nearby SNPs than non-eQTL SVs, which implied that expression-associated SVs tended to occur on local haplotypes that were at least partly represented by SNP markers. This interpretation is consistent with livestock studies showing that the extent to which SVs are tagged by nearby SNPs varies by SV class, allele frequency, SNP panel, and local genomic context (Geibel et al. 2022). However, local SNP tagging did not mean that the SNP-eQTL lead marker captured the same regulatory signal. In conditional regression analyses using SNP-eQTL lead SNPs, SV terms were more often retained than SNP terms, indicating that lead SNPs frequently did not fully represent the expression-associated haplotypes captured by imputed SV dosages. By contrast, conditioning on the highest-LD nearby SNP attenuated many SV associations, as expected if the max-LD SNP provided a stronger local haplotype tag than the SNP-eQTL lead marker. These results indicate that imputed SVs and nearby SNPs often reflect shared local haplotypes, while imputed SV dosages can provide an SV-based representation of regulatory signals that is not always captured by an individual SNP-eQTL lead marker.

This interpretation also provides a useful context for the enrichment of short-read-derived SVs among SV-eQTL candidates. Although short-read-derived SVs represented only a small fraction of the imputed variant set, they contributed disproportionately to expression-associated SVs. This pattern was not explained by an excess of rare alleles among the retained short-read-derived variants, because their allele-frequency distribution was slightly shifted toward higher frequencies rather than toward rare variants. Instead, the enrichment is consistent with discovery-panel ascertainment. This follows a simple sampling principle: the number and frequency distribution of variants observed in a discovery panel depend on both the number of sampled individuals and the underlying allele-frequency spectrum (Cotter et al. 2023). A two-individual long-read discovery panel has limited power to capture variants unless they are relatively common, whereas a larger short-read discovery panel can recover additional segregating variants that are absent from the long-read-derived catalog. The short-read-derived SV-eQTLs therefore illustrate that supplementing a long-read catalog with short-read calls can add functional value, particularly when long-read sequencing is used for discovery in only a small number of individuals. This observation is also relevant to future pangenome-based resources. Although pangenome and graph-based approaches can improve variant representation, our results emphasize that the number and diversity of long-read individuals used for variant discovery will still shape which structural variants are available for downstream genotyping, imputation, and molecular association analyses.

The retained SV-eQTL candidates were not restricted to a single genomic configuration. The most direct configuration was target-gene overlap, illustrated by the PGRMC1-like cis locus, where the associated deletion intersected the target gene body and was annotated with transcript-related consequences. However, other retained cis associations did not directly overlap the target gene. The pias2 locus, for example, represented a nearby cis configuration, while the short-read-detected insertions associated with sparta and slc35a1 occurred near the target genes or within other annotated gene contexts and showed opposite directions of expression effect. These examples indicate that SV-eQTLs cannot be reduced to a simple model in which structural variants disrupt target genes and decrease expression. Previous SV-eQTL studies have shown that structural variants can influence gene expression through several genomic routes, including direct gene overlap, altered copy number, disruption of regulatory sequence, effects on nearby genes, and haplotype-level changes in gene content or regulatory architecture. A recent analysis of the human HLA locus similarly linked complex structural haplotypes to differences in HLA gene expression, highlighting structural variation as a contributor to regulatory diversity at a highly polymorphic immune locus (Canalda-Baltrons et al. 2026). Our retained SV-eQTL candidates fit within this broader pattern, but extend it to an imputation-based setting.

Several limitations define the interpretation of this study. First, SV genotypes in the RNA-seq cohort were imputed rather than directly observed, so SV-eQTL signals depend on reference-panel representation, imputation accuracy, allele frequency, and local LD structure. Second, the SV catalog was incomplete by design: long-read discovery was performed in a small number of individuals, and variants absent from the discovery set or poorly represented in the reference panel could not be evaluated. Third, conditional regression does not establish causal variants when linked regulatory signals coexist. Finally, the expression data were derived from gill tissue, and the associations reported here should not be assumed to generalize across tissues, developmental stages, or environmental contexts.

Despite these limitations, the framework is broadly applicable to systems in which large transcriptomic cohorts are available but direct long-read sequencing or WGS of all individuals is impractical. The challenge is not only the cost of sequencing. Structural variants are also more difficult than SNPs to discover, genotype, phase, impute, and compare across individuals, because they vary in size, breakpoint resolution, allelic representation, and local sequence context (Mills et al. 2011; Jakubosky et al. 2020; Stuart et al. 2026). Many experimental, breeding, ecological, and evolutionary studies have SNP-array genotypes, pedigree or haplotype information, RNA-seq phenotypes, and a smaller WGS reference panel. In such settings, graph-genotyped SVs can be layered onto existing SNP-based cohorts through imputation, enabling structural variants to be tested alongside SNPs in expression association analyses. Future studies could extend this framework by increasing the diversity and size of long-read discovery panels, using pangenome or graph-reference resources, directly validating selected SVs in expression cohorts, and integrating SV-eQTLs with chromatin accessibility, promoter interaction, and trait association data. Overall, graph-genotyped SVs, when imputed into large RNA-seq cohorts, do not replace SNP-based eQTL analysis and should not be interpreted as SNP-independent by default. Instead, they add SV-based information to SNP-defined regulatory signals, allowing a subset of expression-associated haplotypes to be represented as explicit deletions, insertions, or other structural variants where imputation and local haplotype structure support that interpretation.

## Data Availability

Short-read sequencing data for the 906 offspring and 112 parents are available from the European Nucleotide Archive under project accession PRJEB47441. Oxford Nanopore long-read sequencing data for the two individuals used for SV discovery in this study are available under BioProject accession PRJEB113180.

Processed input tables, downstream result tables, and analysis scripts used for the analyses and figures are available on figshare 10.6084/m9.figshare.32567013. All genomic coordinates are based on the Atlantic salmon reference genome Ssal_v3.1 / GCA_905237065.2.

## Acknowledgments

We acknowledge the use of the Orion computing cluster at the Norwegian University of Life Sciences (NMBU). We also acknowledge the internship program between Université de Brest and NMBU for supporting the internship associated with this study.

## Author contributions

Maëlys Chapis: Formal analysis, Investigation, Visualization, Writing – original draft, Writing – review and editing.

Domniki Manousi: Methodology, Formal analysis, Supervision, Writing – original draft, Writing – review and editing.

Célian Diblasi: Formal analysis, Supervision, Visualization, Writing – review and editing. Cathrine Brekke: Formal analysis, Writing – review and editing.

Jun Soung Kwak: Resources, Investigation, Writing – review and editing.

Arturo Vera Ponce De Leon: Resources, Investigation, Writing – review and editing. Mariann Arnyasi: Resources, Investigation, Writing – review and editing.

Randi Fenstad: Resources, Data curation, Writing – review and editing. Solomon Antwi Boison: Resources, Data curation, Writing – review and editing.

Marie Saitou: Conceptualization, Methodology, Formal analysis, Validation, Supervision, Visualization, Writing – original draft, Writing – review and editing.

